# Remodelling of potassium currents underlies arrhythmic action potential prolongation under beta-adrenergic stimulation in hypertrophic cardiomyopathy

**DOI:** 10.1101/2022.03.01.482518

**Authors:** Ruben Doste, Raffaele Coppini, Alfonso Bueno-Orovio

## Abstract

Hypertrophic cardiomyopathy (HCM) patients often present an enhanced arrhythmogenicity that can lead to lethal arrhythmias, especially during exercise. Recent studies have indicated an abnormal response of HCM cardiomyocytes to β-adrenergic receptor stimulation (β-ARS), with prolongation of their action potential rather than shortening. The mechanisms underlying this aberrant response to sympathetic stimulation and its possible proarrhythmic role remain unknown. The aims of this study are to investigate the key ionic mechanisms underlying the HCM abnormal response to β-ARS and the resultant repolarisation abnormalities using human-based experimental and computational methodologies. We integrated and calibrated the latest models of human ventricular electrophysiology and β-ARS using experimental measurements of human adult cardiomyocytes from control and HCM patients. Our major findings include: (1) the developed in silico models of β-ARS capture the behaviour observed in the experimental data, including the aberrant response of HCM cardiomyocytes to β-ARS; (2) the reduced increase of potassium currents under β-ARS was identified as the main mechanism of action potential prolongation in HCM, rather than a more sustained inward calcium current; (3) dispersion of repolarisation between healthy and HCM tissue was increased upon β-ARS, while transmural dispersion in HCM tissue was reduced; (4) models presenting repolarisation abnormalities were characterised by downregulation of the rapid delayed rectifier potassium current and the sodium-potassium pump, while inward currents were upregulated. In conclusion, our results identify causal relationships between the HCM phenotype and its arrhythmogenic response to β-ARS through the downregulation of potassium currents.

## 1. Introduction

Hypertrophic cardiomyopathy (HCM) is the most common inheritable heart pathology and the main cause of sudden cardiac death in young adults. It has a prevalence of 1 in 500 in the general population and is an important cause of arrhythmic sudden death, heart failure, and atrial fibrillation [1]. HCM is usually characterised by left ventricular hypertrophy, which appears in different areas of the ventricle and can lead to left ventricle outflow tract obstruction or, in severe cases, systolic dysfunction and heart failure [2]. However, one of the most dangerous consequences of HCM is the high risk of developing lethal arrhythmias, especially in the young population [3,4]. The specific causes of the high incidence of these lethal arrhythmic events are still not well understood [5,6]. The development of fibrotic tissue in the hypertrophic regions, and consequently the creation of possible re-entry circuits, were originally thought to be the origin of such arrhythmias [7]. However, since most sudden cardiac death cases occur in patients at earlier stages of the disease, when no fibrotic tissue is still present, strong evidence suggests that arrhythmias are not the sole consequence of fibrotic remodelling [5,8].

Conversely, it has been supported that the main factor underlying arrhythmic events in HCM is the altered electrical function of remodelled HCM cardiomyocytes, acting as arrhythmogenic substrates or initiating triggers [9]. Indeed, human HCM cardiomyocytes develop an aberrant electromechanical phenotype, characterised by a prolonged action potential (AP) and a calcium transient (CaT) with slower decay kinetics, as a result of an overexpression of the L-type calcium (I_CaL_) and late sodium (I_NaL_) currents, a downregulation of potassium repolarising currents, and intracellular calcium handling remodelling [9]. The resulting HCM phenotype is more prone to arrhythmia, showing an increase in the number of repolarization abnormalities, such as early afterdepolarisations (EADs) [9,10]. A recent study [10] has also shown that the occurrence of these events is greatly increased under conditions of β-adrenergic receptor stimulation (β-ARS), in parallel with the abnormal electrophysiological response of the HCM cardiomyocytes.

β-ARS is a physiological response mechanism that regulates cardiomyocyte activity, producing a positive inotropic (enhanced contraction), lusitropic (faster relaxation), and chronotropic (increased heart rate) effect. It is triggered by the sympathetic nervous system under either stress or exercise conditions. β-ARS response affects cell membrane ion channels, exchangers, and pumps, and therefore regulates cellular calcium handling and repolarisation. The binding with β-ARs activates the receptor-bound stimulatory G protein, which enhances the production of 3’-5’-cyclic adenosine monophosphate (cAMP), activating protein kinase A (PKA). PKA phosphorylates key cellular substrates that affect the functioning of several channels and ion pumps. Its main sarcolemmal targets are I_CaL_, the slow delayed rectifier potassium (I_Ks_) channels, the fast sodium current (I_Na_) channels, and the sodium-potassium pump (I_Nak_). At the subcellular level, PKA also phosphorylates the ryanodine receptors (RyRs) and phospholamban (PLB), both located on the sarcoplasmic reticulum, as well as troponin I (Tn-I), myosin binding protein-C, and titin, these located on the myofilaments. All the aforementioned phosphorylation processes accentuate the unbalance of ionic currents associated with HCM remodelling, leading to an abnormal response to β-ARS, characterised by a prolongation of the AP duration (APD) and an increase in repolarisation abnormalities [5,10]. Such an abnormal β-ARS response can therefore act as one of the main triggers of arrhythmic episodes in HCM, even in the absence of a fibrotic substrate. From a clinical standpoint, this can be associated to the enhanced arrhythmogenicity and sudden cardiac death of HCM patients during exercise, especially in competitive athletes [11]. It is also reflected in the routine β-blocker therapy in these patients, as well as in current clinical guidelines, recommending reduced exercise intensity or its complete interruption in the most severe cases [2]. However, the main mechanisms underlying the aberrant β-ARS response in HCM remain unknown [5].

We hypothesised that the use of computer models can help understand the pathophysiology and the different interactions between electrophysiology, calcium handling, and β-ARS response in HCM. In recent years, Passini et al. [12] and Coppini et al. [13] have used electrophysiological models to study ion channel remodelling, repolarisation abnormalities, and drug response in HCM cardiomyocytes, while works such as O’Hara et al. [14], Xie et al. [15], or Mora et al. [16] have integrated β-ARS effects in different cardiac pathologies, such as long QT syndrome or heart failure. In particular, in this work we aim to investigate the key ionic mechanisms underlying the HCM abnormal response to β-ARS using human-based experimental and computational methodologies, and to assess how ionic remodelling under β-ARS affects the initiation of repolarisation abnormalities in HCM. To do so, we developed an electrophysiological model that incorporates HCM pathophysiology and β-ARS, based on human experimental data.

We use a population of models approach to isolate the main changes of cellular response in HCM under β-ARS, and the principal ionic mechanisms underlying them, together with assessing the arrhythmogenicity of the models. The developed model and conducted simulation studies offer new insights on the mode of function of β-blockers in HCM and on the ionic mechanisms underlying proarrhythmic events in the disease, as well as on the identification of targets for the development of future anti-arrhythmic treatments.

## 2. Methods

### 2.1 Experimental Data

Electrophysiological data was collected at University of Florence. HCM samples were obtained from a total of 23 HCM patients that underwent surgical myectomy to relieve left ventricle outflow tract obstruction. The control cohort included samples from 8 patients that required a septal myectomy during aortic stenosis or regurgitation surgery and were nonrelated with HCM. The studies were approved by the local ethics committee and written informed consent was obtained from all participants.

Membrane potential was measured using the perforated patch whole-cell current-clamp technique. Calcium transients were registered using the calcium-sensitive fluorescent dye FluoForte. Whole-cell ruptured patch voltage-clamp was used to record I_CaL_ and delayed rectifier K^+^ currents. Details about protocols and solutions can be found in Coppini et al. [9]. Isoproterenol (ISO) at a concentration of 100 nM was used to elicit β-ARS in the myocytes. Test recordings in the presence of the drug were performed after >3 minutes from the beginning of drug exposure. Afterwards, ISO was washed out for >5 minutes and measurements were repeated.

### 2.2 Electrophysiology model

#### 2.2.1 Human baseline electrophysiological model

We developed a mathematical model based on the experimental data that integrates human cellular electrophysiology and β-ARS response. The ToR-ORd model [17] was used to represent human cardiomyocyte electrophysiology. This recently developed model includes an updated formulation for I_CaL_, a critical component in HCM, better recapitulating human experimental recordings [17]. The model also includes updated formulations for the rapid delayed rectifier potassium (I_Kr_), inward rectifier potassium (I_K1_), and I_NaL_ currents. As a result, the AP plateau and the APD adaptation also presented a better agreement with human experimental data than previous electrophysiological models.

#### 2.2.2 Human HCM phenotype

The human HCM phenotype was replicated by scaling the cell volume, ionic conductances, and time constants of the control model, based on the acquired experimental data and the results reported by Coppini et al. [9] and Passini et al. [12]. We also modified I_CaL_ inactivation to reproduce a less functional response of voltage dependant inactivation in HCM, as informed by the experimental data. Specifically, we added a voltage shift in the j_ca_ gate. A detailed description of all these changes is provided in the Supplementary Material.

#### 2.2.3 ϐ-ARS model

The electrophysiological model was combined with the model of β-ARS originally developed by Heijman et al. [18] for canine, and adapted to human electrophysiology by Gong et al. [19]. The model includes β1 and β2 isoforms in β-ARS stimulation, using four conformations to represent the phosphorylation of targets in the PKA and Ca^2+^/calmodulin-dependent protein kinase II (CaMKII) cascades (phosphorylated by PKA, by CaMKII, by both, and non-phosphorylated). A representation of the model is provided in Figure 1, where the different channels and proteins phosphorylated by PKA are represented with an encircled orange *P*, while channels phosphorylated by CaMKII are written in green. In total, the model considers 8 different PKA targets: I_CaL_, I_Ks_, PLB, Tn-I, RyR release, I_NaK_, and fast Na^+^ (I_Na_) and background K^+^ (I_Kb_) currents.

**Figure 1:**
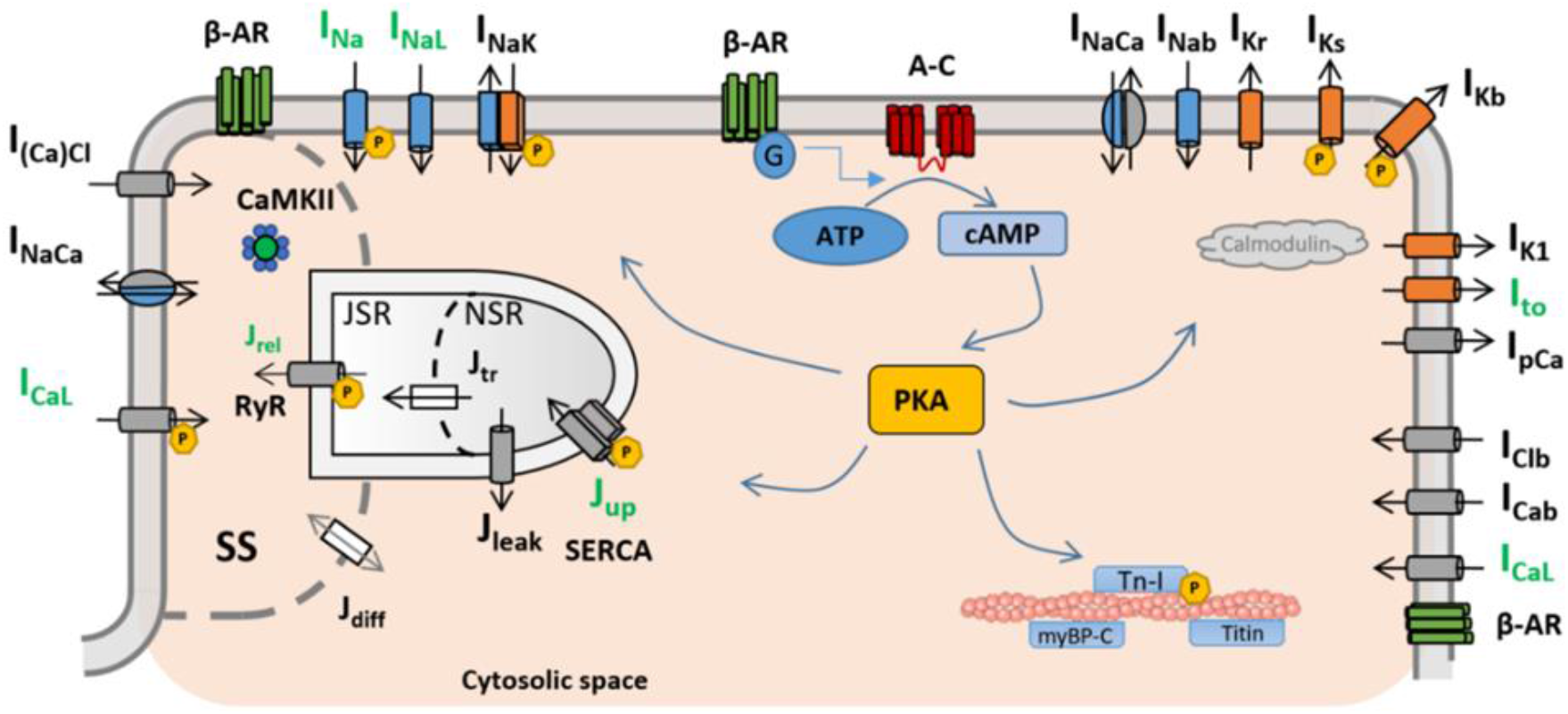
Schematic representation of the ionic currents and subcellular targets considered in the ToR-ORd model integrated with ϐ-ARS response. Currents with proteins phosphorylated by PKA are marked with an encircled P, while CaMKII targets are shown in green. Adapted from [42] under the Creative Commons Attribution (CC BY 4.0) license.

Further modifications were introduced to integrate the human electrophysiological and β-ARS models, and to reproduce the HCM experimental measurements in human cardiomyocytes (see Supplementary Material). Specifically, following the procedure explained by Tomek et al. [17], we implemented a new I_CaL_ activation curve under β-ARS obtained by using a non-linear driving force, and recalibrated the ISO response by reducing I_CaL_ increase under β-ARS (I_CaL_ augmented by 10% instead of 40%, compared to [19]). This modification was required due to the distinct formulation of the ToR-ORd model, which presents higher baseline I_CaL_ values. Following experimental results in HCM cells, ISO-induced changes in the amplitudes of I_CaL_ and I_Ks_ and in I_CaL_ inactivation were also included (see section 3.1). The code (Matlab) for the simulation of control and HCM models under β-ARS is available at https://github.com/rdoste.

#### 2.2.4 Populations of models and simulation protocols

An initial population of 2,000 human AP and CaT control models was built applying parameter variability, generating different sets of conductances for the main channels and pumps (I_CaL_, I_Na_, I_NaL_, I_to_, I_Ks_, I_Kr_, I_K1_, I_NaCa_, I_NaK_, J_rel_, J_up_) via Latin hypercube sampling, as described in Britton et al. [20] and Passini et al. [12]. All parameters were sampled in the (50%-150%) range with respect to the original baseline values. The control population was calibrated according to the ranges extracted from human experimental data and detailed in [12]. After calibration, 573 models were retained in our final control population. The HCM population was built from the control one by applying the previously described ionic remodelling.

Cells were stimulated with a depolarising current of −53 pA/pF during 1 ms [17]. An ISO concentration of 100 nM was used as an input of the simulations. Models were stimulated at 1 Hz for 350 s to ensure a steady state in the phosphorylation of the PKA targets. All the simulated and experimental results are reported as mean±SD.

### 3. Results

#### 3.1 Model calibration with experimental data recapitulates the abnormal electrophysiological response of human HCM cardiomyocytes under β-ARS

The altered electrophysiological response of HCM cells in both baseline and β-ARS conditions was reproduced by the proposed computational model by incorporating ionic remodelling in different channels and intracellular calcium fluxes (see Methods), according to the experimental findings. Selected experimental results of control and HCM cardiomyocytes at 1 Hz pacing are presented in Figure 2. β-ARS response by ISO resulted in APD shortening at 90% of repolarisation (APD_90_) by −16±5% in control, while it aberrantly prolonged by 12±3% in HCM cardiomyocytes. The amplitude of the CaT in HCM was increased by 55±14%, while calcium transient decay at 50% relaxation (T_50_) was accelerated (−28±11%). L-type calcium current magnitude increased to a similar extent in both cell types during β-ARS (+34±12% and +30±10%, control vs HCM). The duration of the I_CaL_ current (from depolarisation onset to 50% of decay) was also measured. Under β-ARS, I_CaL_ in control cells presented approximately the same duration (+3±6%), while in HCM cells the duration increased by 22±6%. Finally, experimental data also showed an abnormal β-ARS response in potassium currents. Total I_k_ density increased by 69±10% in control cells during ISO application. However, I_K_ density in HCM cardiomyocytes only increased by a 39±7%.

**Figure 2:**
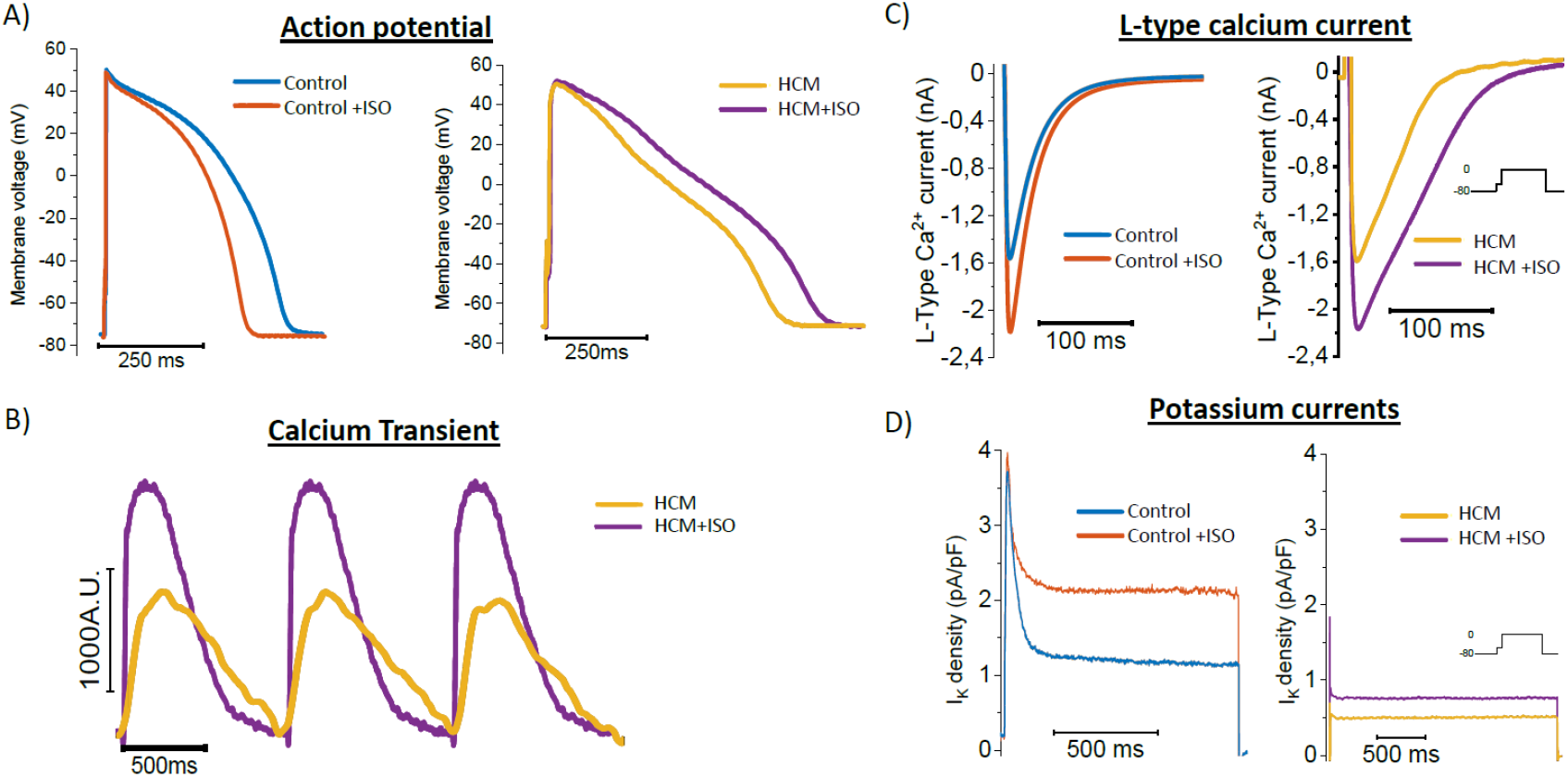
Experimental recordings of action potential (A), calcium transient (B), L-type calcium current (C), and potassium currents (D), for selected control (blue) and HCM (yellow) human samples. Recordings obtained after ϐ-ARS response by isoproterenol (ISO) are superimposed in orange (control) and purple (HCM).

I_CaL_ and potassium currents of the HCM cardiomyocyte model were calibrated based on the human experimental findings under β-ARS, leading to similar AP and CaT response. As detailed in the Supplementary Material, I_CaL_ density was increased by 10% under β-ARS, which together with the implemented changes in activation, led to a similar increase in I_CaL_ magnitude as in the experimental findings (see later Figure 4). Its fast and slow inactivation constants were also doubled, to mimic the more persistent I_CaL_ recorded under β-ARS in HCM cardiomyocytes. Finally, the increase of the I_Ks_ and I_Kb_ currents was reduced by 90%, considering basal differences in the expression of these currents. As the experimental recordings could not measure independently the different components of the total potassium current due to its small amplitude in HCM, we only applied this modification to previously reported potassium channels affected by the β-ARS response, i.e., I_Ks_ and I_Kb_.

Figure 3 shows the reproduction of the β-ARS response in the simulated populations of 573 control and HCM cellular models. A representative case of AP behaviour is presented in Fig. 3A, which shows how the AP prolongation obtained in the HCM phenotype under β-ARS (in purple) differs from the reported AP shortening in control cells (orange). Calculated APDs for both populations and protocols are displayed in Fig. 3B, showing that the abnormal AP prolongation in HCM is a common characteristic to all the simulated HCM models. Similarly, representative CaTs in HCM are presented in Fig. 3C. Biomarkers of CaT amplitude and decay (T_90_ and T_50_) are shown in Figs. 3D-3F. Following the experimental data presented here and in [9,10,12], CaT amplitude in the absence of β-ARS was decreased in HCM, also presenting a slower decay. Under β-ARS, CaT amplitude increased to a larger extent in HCM, while both cell types exhibited a faster decay. A more detailed comparison of the CaT between representative control and HCM models is provided in the Supplementary Materials (Figure S1). Models manifesting any repolarisation abnormalities were not included in our report of AP and CaT statistics.

**Figure 3:**
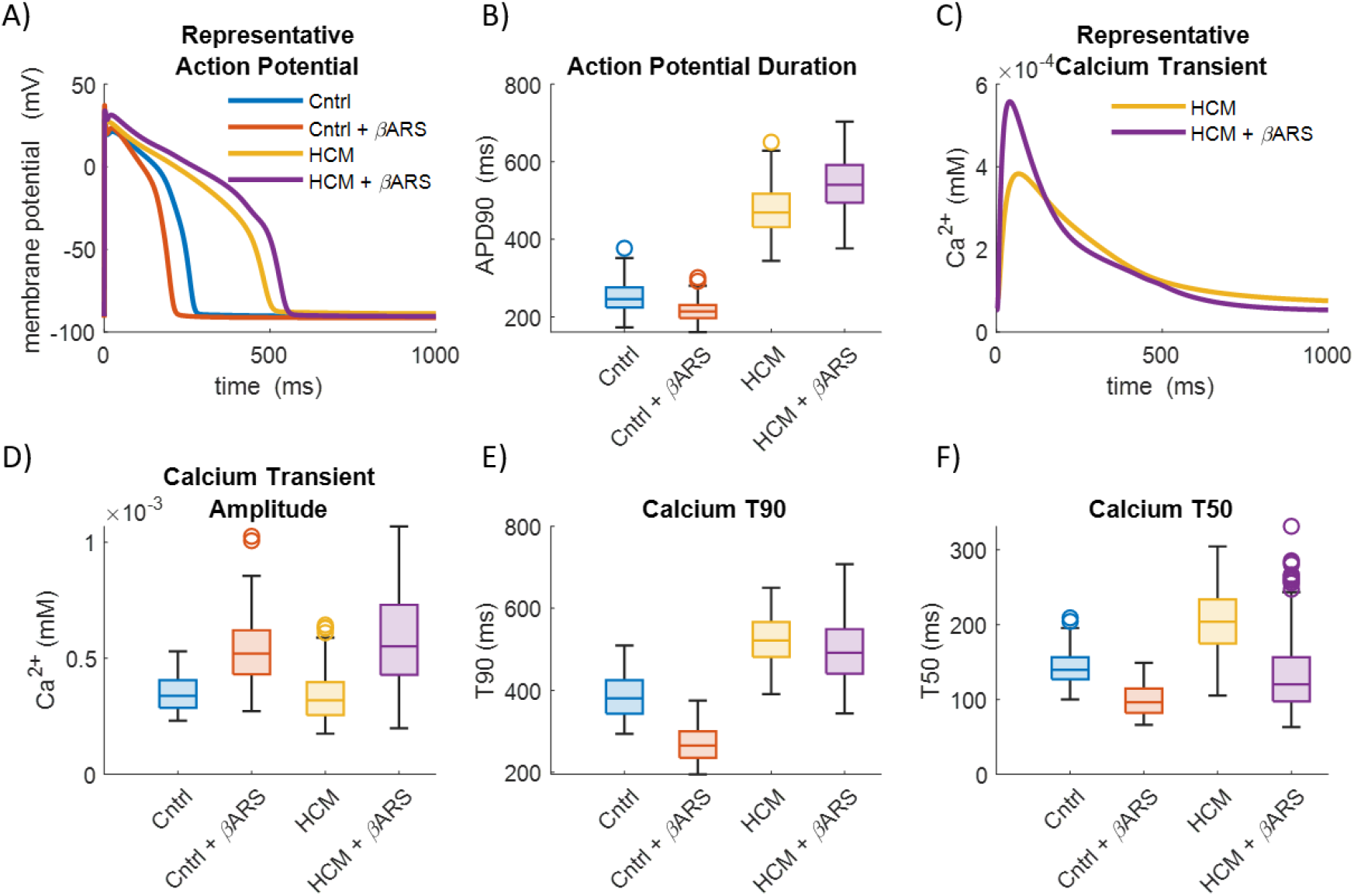
Effect of ϐ-ARS on simulated action potential (AP) and calcium transients (CaT) in control and HCM populations. A) Comparison of the AP of baseline models. B) AP duration of all simulated models. C) Comparison of the CaT of the HCM baseline models. D) CaT amplitude of all simulated models. E) CaT relaxation time from peak at 90% of the decay (T90). F) CaT relaxation time from peak at 50% of the decay. Box outlines show the 25^th^ and 75^th^ percentiles, median is represented by the middle line, and whiskers indicate the 5^th^ and 95^th^ percentiles.

Changes in simulated APs and in CaTs were validated against experimental data. In control, both data presented a similar APD reduction (−13±7% vs −16±5%, simulated vs experimental), while in HCM both showed AP prolongation (14±4% vs 12±3%). CaTs in HCM also presented an increase in the amplitude (73±23% vs 55±14%, simulated vs experimental) and faster decay values of T_50_ (−36±11% vs −28±11%).

#### 3.2 I_Ks_ remodelling underlies the aberrant AP prolongation in HCM under β-ARS

The most significant ionic currents underlying the AP waveform in our baseline control and HCM models are depicted in Fig. 4, allowing the identification of causal mechanisms in the divergence to β-ARS response between control and HCM cardiomyocytes. In particular, I_CaL_ exhibits a slower inactivation in HCM, which is even slower under β-ARS. Nevertheless, the biggest differences are observed in I_Ks_ under β-ARS, increasing to a much smaller extent in HCM than in control. HCM cells also present higher baseline values of I_NaK_ and I_NaL_, which are additionally increased under β-ARS. β-ARS also rose the amplitude of I_Na_ currents and J_rel_ and J_up_ fluxes. In the HCM case, J_up_ flux also showed slower kinetics.

**Figure 4:**
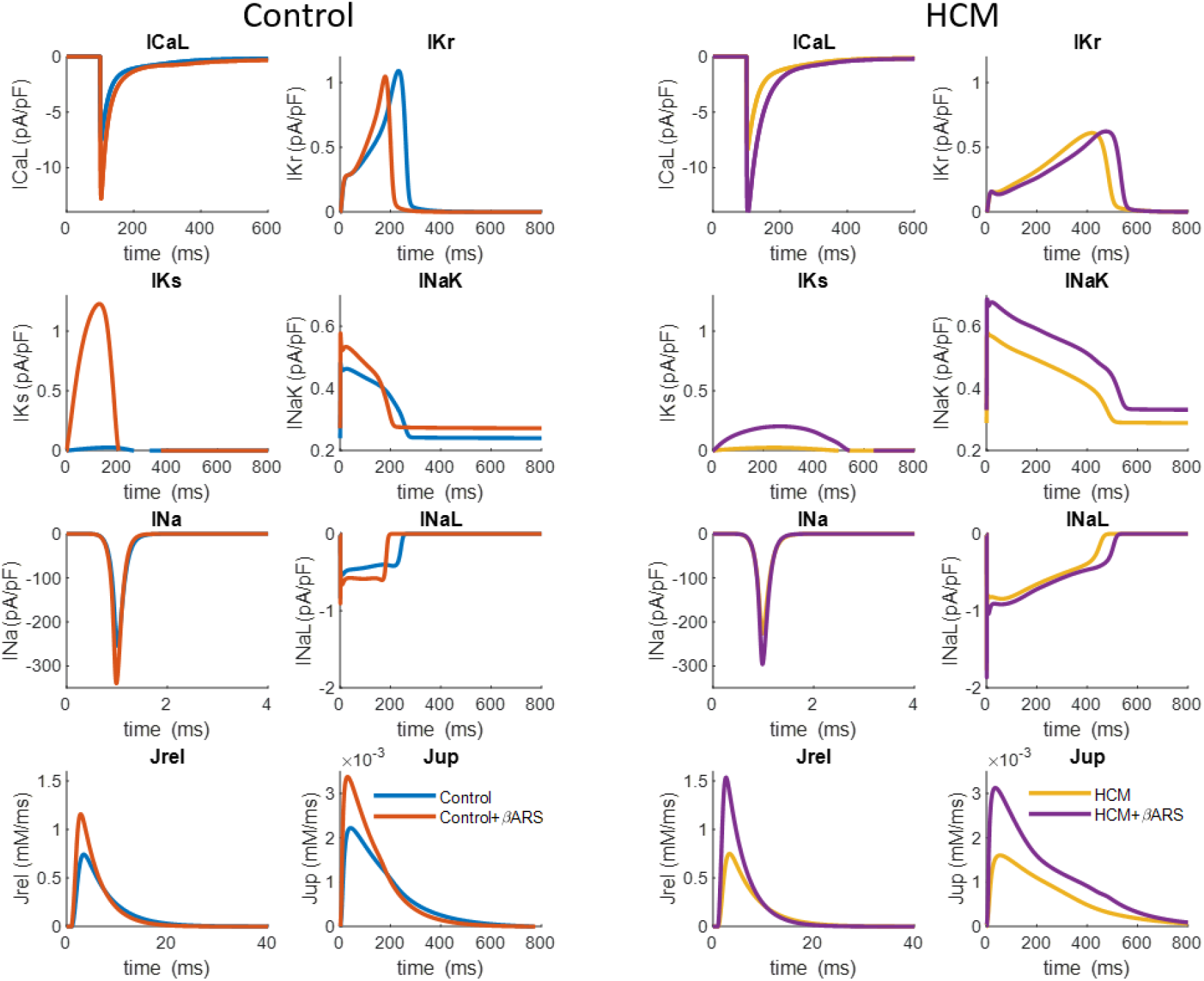
Effect of ϐ-ARS on ionic currents for control (left columns) and HCM (right columns) baseline models. The different represented traces are: L-Type calcium current (I_CaL_), rapid delayed rectifier potassium current (I_Kr_), slow delayed rectifier potassium current (I_Ks_), sodium-potassium pump (I_NaK_), fast sodium current (I_Na_), late sodium current (I_NaL_), the calcium RyR release flux (J_rel_), and calcium SERCA pump uptake flux (J_up_).

In order to determine the main mechanisms underlying the abnormal AP prolongation in HCM under β-ARS, we performed a sensitivity analysis of the main phosphorylation targets to quantify the individual role of each current on APD and systolic CaT (CaT_max_). Figure 5 shows the percentage change that each of the PKA phosphorylation targets causes to the APD_90_ and CaT_max_ biomarkers when they are phosphorylated individually. For these experiments, phosphorylation fractions of the rest of the individual targets were kept to 0.

**Figure 5:**
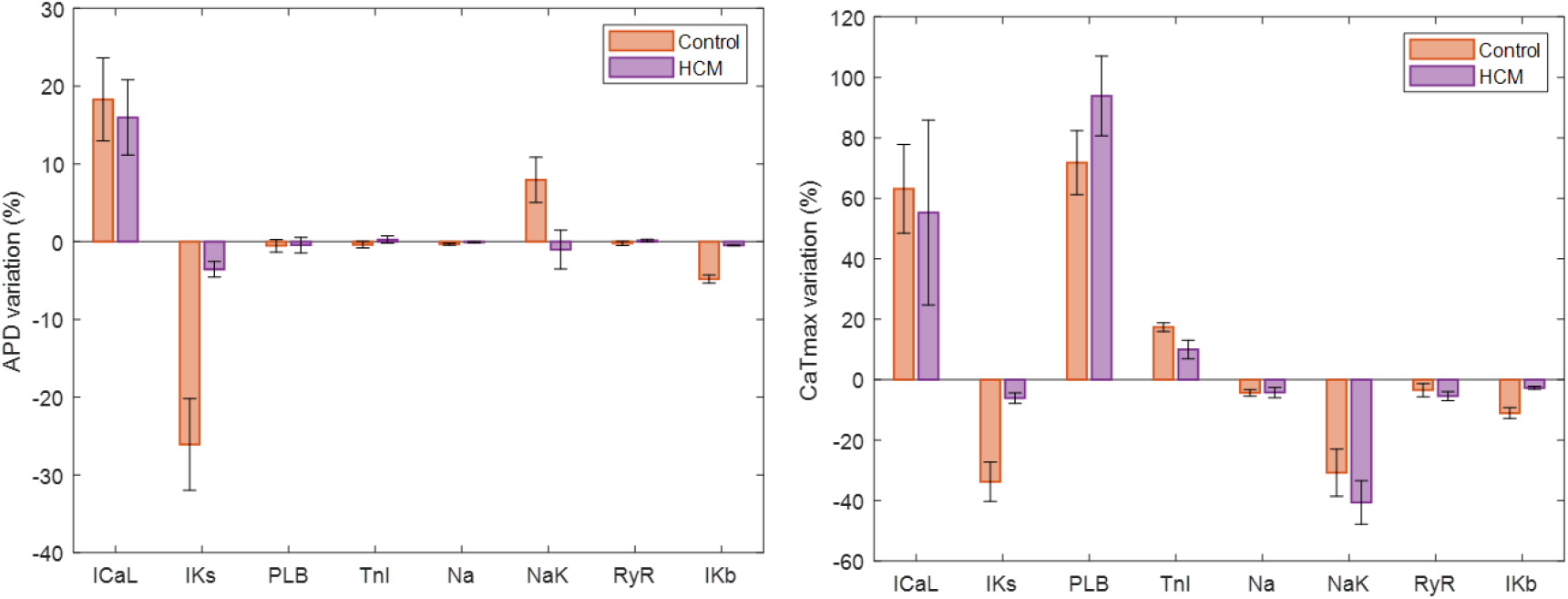
Sensitivity analysis of the contribution of each phosphorylated target to APD_90_ (A) and CaT_max_ (B), calculated for control (red) and HCM (purple) models.

The main modulators of APD under β-ARS in the healthy control models are I_CaL_ and I_Ks_, whose combined contribution leads to AP shortening. However, I_Ks_ impact on APD and CaT is drastically reduced in HCM cases (APD_90_ decreases from −26±6 % in control to only −3.5±1 % on HCM, while CaT_max_ decreases from −33±6% to −6.1±2%). I_Kb_ contributions to both biomarkers are also significantly reduced, although their impact on APD_90_ and CaT is minor. Importantly, the slower inactivation and increased amplitude change of I_CaL_ does not affect the changes in APD or CaT amplitude as much as the reduced augmentation of I_Ks_. Taken together, our results identify the experimentally reported reduction of I_K_ currents augmentation under β-ARS as the main difference in the response to β-ARS between control and HCM cardiomyocytes, becoming insufficient in HCM cells to reduce the APD and CaT in a similar extent than in control cells.

#### 3.3 Dispersion of repolarisation increases in HCM myocardium under β-adrenergic stimulation

Dispersion of repolarisation is associated with an increase in the arrhythmogenicity and can be affected by β-ARS response. In addition to transmural dispersion of repolarisation, HCM is also characterised by regional mosaicism, where only hypertrophied regions may exhibit ion channel remodelling, as main explanatory effect of ECG abnormalities in T-wave morphology in the disease [13,21], making it important to also investigate dispersion between HCM-remodelled areas and normal myocardium. We therefore simulated the effects of β-ARS in the APs of both HCM and healthy myocytes, including endocardial and epicardial cells, the latter of especial importance as epicardial biopsies are difficult to obtain from in-vivo patients.

Simulation results of APD changes in endocardial vs epicardial cells, and their percentage of APD variation, are presented in Figs. 6A-6B. Our findings indicate that both cell types exhibit a similar reduction of APD_90_ under β-ARS in control (−13±7% vs −15±7%, respectively). In HCM myocardium, however, the extent of APD prolongation was significantly larger in epicardial cells (14±4% vs 23±8%, Fig 6B). This was determined by differences in ion channel expression between both layers [17,22]. More specifically, the larger prolongation was caused by the more predominant epicardial I_CaL_ current, despite its also larger I_Ks_ density, due to the reduced augmentation of potassium currents in the HCM response to β-ARS.

**Figure 6:**
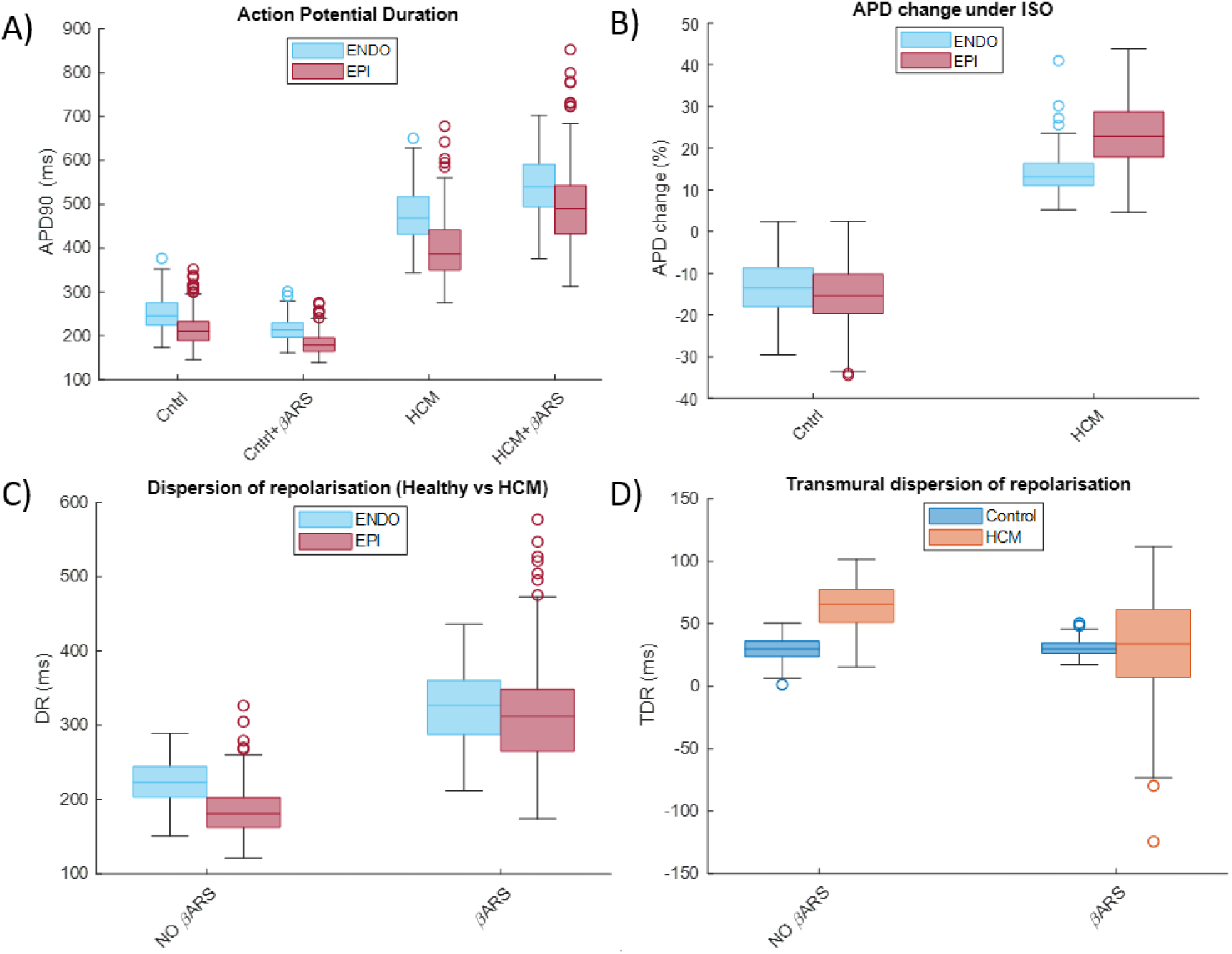
Effect of ϐ-ARS stimulation on APD_90_ and dispersion of repolarisation. A) APD_90_ distribution for endocardial (ENDO) and epicardial (EPI) populations, in control and HCM myocytes, in basal conditions and under the application of ISO. B) Percentage change in APD_90_ in both cell types after ϐ-ARS stimulation by ISO. C) Dispersion of repolarisation between control and HCM-remodelled cells. D) Transmural dispersion of repolarisation in control and HCM tissue, before and after ϐ-ARS stimulation by ISO. Box outlines show the 25^th^ and 75^th^ percentiles, median is represented by the middle line, and whiskers indicate the 5^th^ and 95^th^ percentiles of the distributions.

The AP prolongation in HCM under β-ARS stimulation, common to both the endocardial and epicardial cells, resulted in an increased dispersion of repolarisation (DR=APD_90,HCM_-APD_90,control_) between the healthy and HCM remodelled tissues: from 223±27 ms in baseline to 324±47 ms under ISO in the endocardium, and from 186±34 ms in baseline to 314±69 ms under ISO in the epicardial layer (Fig. 6C). Transmural dispersion of repolarisation (TDR=APD_90,ENDO_-APD_90,EPI_) was also studied (Fig. 6D). Application of ISO did not induce any change in control cells; however, the higher transmural dispersion of repolarisation in HCM decreases after β-ARS (from 64±17 in baseline to 32±37 ms under ISO), due to the higher prolongation of epicardial cells with respect to the endocardial ones. Nevertheless, the dispersion of repolarisation between control and HCM myocardium increased to a much larger extent than the observed transmural reduction, which can contribute to the increased arrhythmic risk in HCM patients under β-ARS stimulation.

#### 3.4 Faster I_CaL_ phosphorylation kinetics underlie EADs formation under β-ARS in HCM

We finally studied the mechanisms underlying the development of EADs in HCM under β-ARS. In simulated HCM cardiomyocytes, repolarisation abnormalities were detected in 273 out of 573 models in endocardial cells, and in 308 out of 573 in their paired epicardial configurations. EADs were rarely observed in simulated control myocytes (3 out of 573 models). Simulated HCM AP traces are presented in Fig. 7A. Figure 7B shows the distributions of ionic properties for the HCM models that developed EADs under ISO application, compared to those exhibiting a regular repolarisation phase. All models that presented repolarisation abnormalities were characterised by an unbalance of upregulated inward (I_CaL_, I_NaCa_, and I_NaL_) and downregulated outward (I_kr_ and I_NaK_) currents.

**Figure 7:**
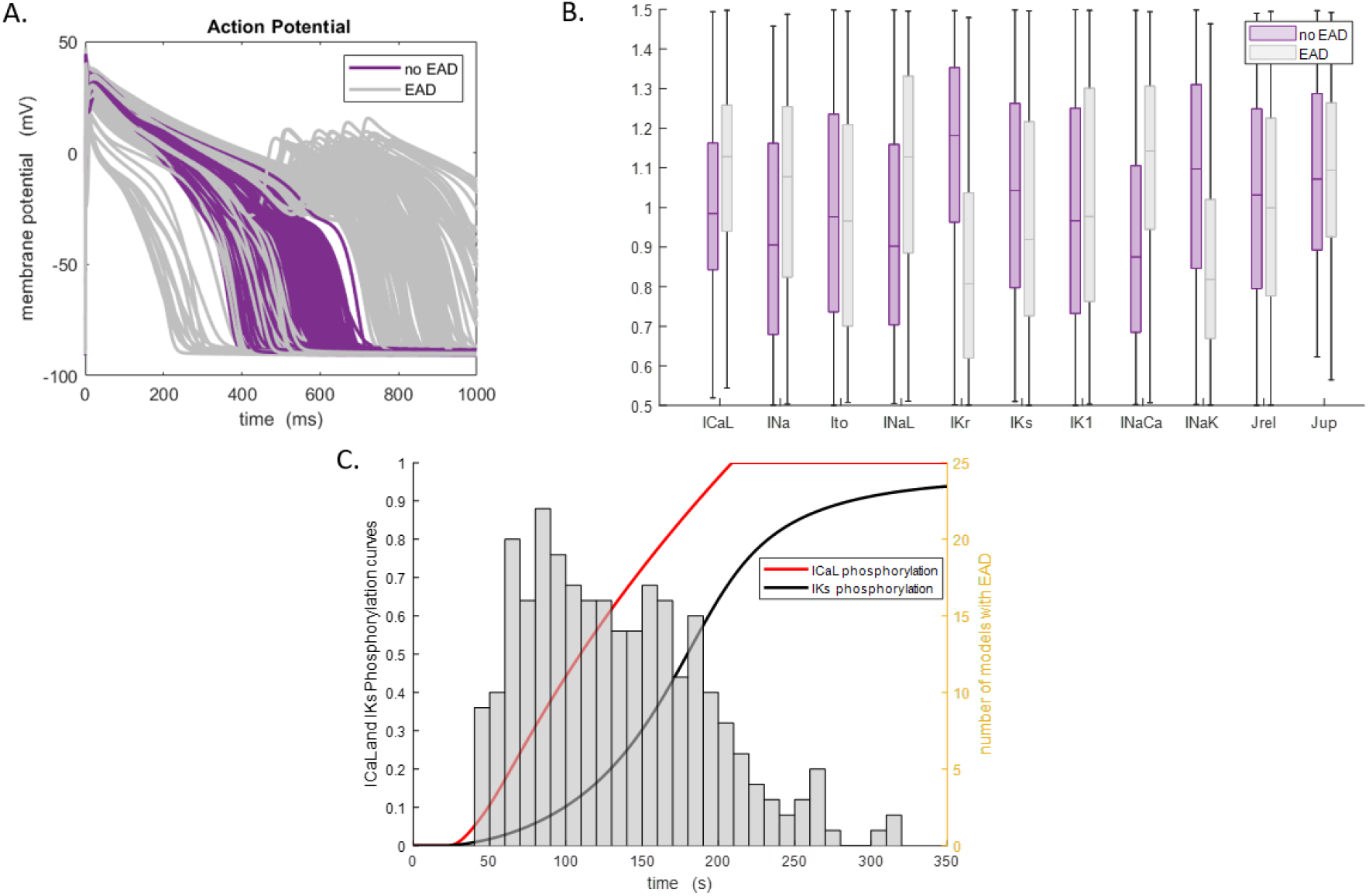
Effect of ϐ-ARS stimulation on EADs onset in HCM. A) AP traces of HCM models not developing (purple) and developing (grey) EADs under ϐ-ARS in HCM endocardial cells. B) Normalised distribution of scaling factors of ion channel conductances for same models as in A). Box outlines show the 25^th^ and 75^th^ percentiles, median is represented by the middle line, and whiskers indicate the 5^th^ and 95^th^ percentiles. C) Number of models that developed their first EAD at a certain interval of time. The frequency histogram is superposed with the level of phosphorylation of I_CaL_ and I_Ks_ currents (ranging from 0 to 1).

To further study the mechanisms causing repolarisation abnormalities under β-ARS, we analysed the time of first EADs occurrence against the phosphorylation of targets by β-ARS. Figure 7C illustrates the number of models developing EADs against the time of the first incidence, together with the evolution over time of the phosphorylation percentage for I_CaL_ and I_Ks_ currents (0 and 1 respectively meaning no and full phosphorylation). ISO administration is simulated after 20 s. Results show that the majority of the EADs start just after the phosphorylation cascade initiates, between 50 to 100 s, coinciding with the maximal difference between I_CaL_ and I_Ks_ phosphorylation curves. This suggests the much faster phosphorylation of I_CaL_ compared to a weakened I_Ks_, therefore unable to counter-balance the total inward current increase, as the main precursor of cellular repolarisation abnormalities under β-ARS in HCM.

## 4. Discussion

Myocyte remodelling under β-ARS stimulation, as observed in HCM patients, directly affects the electrophysiological response of the heart and is strongly related to arrhythmic episodes, especially under exercise conditions. In this study, we developed a human HCM ventricular electrophysiological model with an integrated β-ARS response, based on experimental recordings from human healthy and HCM cardiomyocytes. The main findings of the present human-based experimental and computational study are: (i) the developed model was able to replicate the HCM phenotype and its behaviour under β-ARS, including abnormal AP prolongation and increase in the propensity to arrhythmia development; (ii) remodelling of potassium channels was found as the main factor underlying the abnormal AP prolongation of HCM cardiomyocytes under β-ARS; (iii) simulation studies of endocardial and epicardial HCM phenotypes showed that transmural dispersion of repolarisation is higher in HCM remodelled tissue, but it is reduced under β-ARS. Conversely, β-ARS stimulation increases the dispersion of repolarisation between healthy and HCM myocardium, both in the epicardial and endocardial layers; (iv) the faster phosphorylation of I_CaL_ underlies EADs formation in HCM cardiomyocytes under β-ARS. Such proarrhythmic events occurred in HCM cardiomyocytes exhibiting upregulated I_CaL_ and I_NaL_ and downregulated I_Ks_ and I_NaK_ currents, being these currents the main targets for future pharmacological treatment.

### 4.1 Development of a β-ARS model for human HCM phenotype

The human electrophysiological studies conducted in this work were enabled by developing a coupled version of the ToR-ORd [17] with the Heijman β-ARS phosphorylation [18] models. Although other works [14,19] have adapted the Heijman model to human models [22], this work is the first to adapt it to the ToR-ORd model formulation. The ToR-ORd model represents an improved description of human cellular electrophysiology than its predecessors, with a better description of the human AP plateau, APD accommodation, and calcium dynamics. The model also includes an updated formulation of the I_CaL_ current, which has a key role in β-ARS response and HCM remodelling, and is also one of the main contributors to the abnormal response of HCM cardiomyocytes to β-ARS according to our experimental findings. The model has also proven suitable for replication of arrhythmia precursors, such as EADs and alternans [17,23].

Model adaptation to reproduce with accuracy experimental data was not straightforward. The different formulation of the ToR-ORd model (especially the I_CaL_ current) required additional changes in current conductances, activation curves, and time constants. All of them are described in the methodology section and Supplementary Material. I_CaL_ density increase under β-ARS was reduced compared to previous works, in accordance with our experimental data in human (compared to animal) experiments, and also due to the larger I_CaL_ amplitude compared to other models. However, I_Ks_ contribution under β-ARS was considerably increased with respect to the suggested values by the experimental values. This augmentation of the I_Ks_ is consistent with channel damage during isolation and the findings of several previous works [24–26], which describe a change of the role of I_Kr_ and I_Ks_ under β-ARS stimulation. I_Ks_ magnitude becomes more important during repolarisation, while I_Kr_, dominant under basal conditions, is barely modified. This behaviour can be observed in the simulated results of Fig. 4.

It has been hypothesised that the slower inactivation of I_CaL_ under β-ARS in HCM cardiomyocytes could be a consequence of an abnormal calcium dependant inactivation (CDI) [10]. Under basal conditions, both CDI and voltage dependant inactivation limit the amount of calcium influx during the AP. Under β-ARS, the CDI is predominant [27]. As detailed in the Supplementary Material, we accounted for these factors by readjusting the j_ca_ gate, which regulates CDI [22], in the HCM electrophysiological model. Shall these changes in CDI are not applied, the resulting simulated HCM models almost ubiquitously present repolarisation abnormalities. This finding further suggests the slower inactivation of I_CaL_ to be related to an abnormal CDI in HCM; otherwise, such long AP characteristics of the HCM cardiomyocytes under β-ARS would not be possible with a normal CDI.

We used the population of models approach [12,19,20] to enable the study of the effect of inter-subject variability, and therefore of possible different responses caused by pathology. The simulated healthy control populations qualitatively and quantitatively reproduced the measured experimental data, mimicking the behaviour of the AP under basal and β-ARS conditions, and showing AP shortening under β-ARS (−16±5% vs −13±7% change in experimental vs simulated results). Despite lacking of experimental control CaT data at 1 Hz due to sample availability, the simulated data agrees with previous works [28], showing an increase in amplitude and a shortening of the CaT relaxation (inotropic and lusitropic response). A similar response has been also reported in previous human β-ARS models [14,19]. The HCM phenotype was also recovered and maintained during β-ARS response. In baseline, APDs were longer than in the control populations, together with a small decrease of CaT amplitude and slower kinetics, as supported by the human experimental data. After ISO application, APD was abnormally prolonged (12±3% vs. 14±4% change, experimental vs simulated results), while CaT behaviour showed an increase in amplitude and a relaxation shortening, maintaining the inotropic and lusitropic response. This experimental behaviour, as also reported in previous ex-vivo studies [5,10], could not be reproduced by the previously existing β-ARS models.

### 4.2 Mechanisms underlying the abnormal AP prolongation in HCM under β-ARS

In healthy human cardiomyocytes, APD changes under β-ARS response are mainly caused by the enhancement of I_CaL_ and I_Ks_ currents [29]. Both currents are augmented under β-ARS, but the repolarising potassium currents prevail over I_CaL_ augmentation, leading to a net APD reduction. However, HCM myocytes exhibit an opposing behaviour, showing AP prolongation under β-ARS. Analysis of the experimental data indicated two possible mechanisms contributing to this phenomenon: either an augmented L-type calcium current, with protracted inactivation kinetics in HCM compared to control, or a lower increase of potassium currents in HCM under β-ARS.

In order to identify what components of HCM remodelled electrophysiology determine such an aberrant response to β-ARS, we performed a throughout sensitivity analysis of independent phosphorylation targets. Our results show that the reduced I_Ks_ augmentation associated with HCM remodelling affected its ability to shorten the APD. Although the prolongation of I_CaL_ in HCM cardiomyocytes favoured the prolongation of the APD, it was not sufficient *per se* to induce the abnormal prolongation, having a similar contribution to APD prolongation as I_CaL_ in control cardiomyocytes. Therefore, our results indicate the reduced augmentation of potassium currents as the key mechanism underlying the abnormal prolongation of the AP in human HCM cardiomyocytes under β-ARS.

In addition, the β-ARS insults affected the inotropic response. The reduced augmentation of I_Ks_ affected positively the CaT amplitude, although this was more determined by changes in the amplitude of calcium currents and intracellular fluxes, such as I_CaL_ and SERCA/PLB. The more sustained I_CaL_ of HCM cardiomyocytes under β-ARS did not significantly affect CaT amplitude, but prolonged its duration.

### 4.3 Modulation of dispersion of repolarisation by β-ARS in HCM

Dispersion of repolarisation is associated with the establishment of arrhythmogenic substrates [30,31]. In healthy hearts, the dispersion of repolarisation is small and usually occurs in the transmural direction, since epicardial cells present shorter APDs than the endocardial ones. Previous studies have shown that β-ARS maintains the transmural dispersion of repolarisation in healthy hearts, but it can greatly increase it in some cardiac pathologies, such as Long QT syndrome [26] or heart failure [28]. Here we studied the modulation of dispersion of repolarisation by β-ARS when combined with HCM remodelled tissue.

According to our results and consistent with previous findings, transmural dispersion of repolarisation was not affected by β-ARS in healthy tissue (Fig. 6D). Conversely, it decreased in HCM myocardium (Fig. 6D), due to the epicardial HCM cells having a greater APD prolongation under β-ARS than the endocardial ones (Fig. 6B). However, the dispersion of repolarisation was drastically increased after ISO application between the healthy and remodelled tissues, both in the epicardial and endocardial layers (Fig. 6C). Such border zones between the remodelled and healthy tissue in HCM could therefore act as areas promoting arrhythmogenicity in HCM patients, especially after β-ARS stimulation (i.e., stress/exercise conditions [1,32]).

### 4.4 Genesis of repolarisation abnormalities in HCM under β-ARS

Long APD is related to an increase in the incidence of EADs [33]. Therefore, the abnormal APD prolongation of HCM cardiomyocytes under β-ARS can also act as a trigger of arrhythmogenic events. In particular, several studies have reported the incidence of such events after β-ARS stimulation [34–36].

Our results offer insights into how HCM remodelling and its interaction with β-ARS can promote EADs. The ionic profile of the models developing EADs under β-ARS was characterised by upregulation of I_CaL_, I_NaL_ and I_NaCa_ and downregulation of I_Kr_ and I_NaK_ currents. Overall, the decrease of the potassium currents and the increase of inward currents during β-ARS is the cause of the occurrence of such repolarisation abnormalities. Confirming previous findings [9,37,38], the main mechanism underlying EADs onset was I_CaL_ reactivation secondary to AP prolongation. Pharmacological strategies targeting overexpressed currents driving AP prolongation in the HCM remodelled phenotype, such as the use of ranolazine [9,39] or disopyramide [13], should be considered for their prevention, with additional benefits over calcium blockers in terms of not compromising CaT amplitude, and therefore the inotropic and lusitropic response.

In our work, we have also explored the kinetics of EADs formation. Several studies [40–42] have indicated different kinetics in PKA phosphorylation targets as modulators of EADs formation, contributing to the establishment of a dynamically impaired repolarisation reserve. In particular, we registered the time of first EAD occurrence in each of our HCM models and studied their dependence with I_CaL_ and I_Ks_ phosphorylation. Our results confirm that the majority of EADs occur when the mismatch between the phosphorylation curves increase, and once reached its maximum difference the occurrence of EADs clearly diminish, confirming the different phosphorylation rate of both targets as a proarrhythmic factor in HCM. Therefore, the development of therapies that could compensate the rates of the different PKA phosphorylation targets could also a represent a promising treatment for these cases of arrhythmia.

### 4.5 Limitations

Although as the first model of its kind directly informed by human data on I_CaL_ and potassium currents in health and disease states, our developed model of coupled human electrophysiology and β-ARS response still remains constrained by the scarcity of human electrophysiological β-ARS data. β-ARS models are largely based on data from other species, reparametrized to human cardiomyocytes. Therefore, differences on distinct ion channel conformations [43], different molecular processes involved in β-ARS [25,44], or distinct densities and locations of β-ARS receptors [45], require further investigations. Data availability regarding myosin binding protein-C and titin phosphorylation [46] would enable implementing these targets in the β-ARS model, qualifying it for quantitative investigations of β-ARS response in active tension. Moreover, human heterogeneity in the distribution of the sympathetic innervation system [47], which could affect epicardial and endocardial differences, was not incorporated into the model.

Human HCM cardiomyocyte data is also scarce, and typically only available from obstructive patients undergoing surgical myectomy. Consequently, the studied HCM phenotype better represents that from patients at later remodelling stages of the disease. The reduced number of samples also limited the number of experimental measurements, easier to perform at slow pacing frequencies to limit myocyte damage. Nevertheless, we based our model and validation using experimental data at 1 Hz, within the normal pacing physiological rate range. Limitations in the measurement of potassium currents, difficult to isolate experimentally, also influenced the implementation of these changes into the model.

### 4.6 Conclusions

We have developed and validated a computational model to study the interplay between β-ARS stimulation and ion channel remodelling in human HCM. The model quantitatively and qualitatively replicated the main electrophysiological alterations observed in human samples, and predicted the reduction of the potentiation of potassium repolarising currents under β-ARS as the main mechanism underlying the aberrant APD prolongation observed in HCM cardiomyocytes under β-ARS. Downregulation of potassium currents was also characteristic of the models that developed EADs, reinforcing a causal relationship between the HCM phenotype and its arrhythmogenic response to β-ARS.

## Acknowledgements

This research was funded by a British Heart Foundation (BHF) Intermediate Basic Science Fellowship to ABO (FS/17/22/32644). RC acknowledges support from the European Union’s Horizon 2020 Research and Innovation Programme under Grant Agreement no. 777204. The authors acknowledge additional support from an Infrastructure for Impact Award from the National Centre for the Replacement, Refinement and Reduction of Animals in Research (NC/P001076/1), and the Oxford BHF Centre of Research Excellence (RE/13/1/30181), PRACE for awarding us access to Piz Daint at the Swiss National Supercomputing Centre, Switzerland (ICEI-PRACE grants icp005 and icp013), and the use of the University of Oxford Advanced Research Computing (ARC) facility (http://dx.doi.org/10.5281/zenodo.22558)

