## Supplementary material for "Remodelling of potassium currents underlies arrhythmic action potential prolongation under beta-adrenergic stimulation in hypertrophic cardiomyopathy"

**EXTENDED METHODS**

### Modifications to the ToR-ORd human ventricular model (control population)

Simulations of healthy myocytes were performed using the last version of the ToR-ORd model [1], with no additional changes except the addition of the dynamic representation of intracellular chloride reported in a posterior addendum [2].

### Modifications to the ToR-ORd human ventricular model (HCM population)

Hypertrophic cardiomyopathy (HCM) phenotype was constructed following the procedure described by Passini et al. [3]. Based on experimental observations [4], we up-regulated $I_{NaL}$ (+165%) and $I_{CaL}$ (+10%), and down-regulated the transient outward potassium ($I_{to}$, -70%) and $I_{K1}$ (-30%) currents; the time constant of fast $I_{CaL}$ inactivation (voltage and calcium-driven) was increased by 35%, while its time constant of slow inactivation was increased by 20%. Based on mRNA expression, we down-regulated $I_{Kr}$ and $I_{Ks}$ currents (-45%), SERCA pump ($J_{up}$, -35%), and RyR release ($J_{rel}$, -20%), while $Na^+/Ca^{2+}$ exchanger ($I_{NaCa}$) was increased (+30%). Following [3], we included an increased affinity of troponin for $Ca^{2+}$ (Ktrpn, -50%), $I_{NaK}$ inhibition (-30%), and an increased background $Na^+$ current ($I_{Nab}$, +165%). Cell radius was also modified to reproduce the 90% increase in cell volume that is present in HCM cells. A comparison of these changes versus the ones performed in the original HCM model, that used the O'Hara-Rudy (ORd) formulation [5], is presented in Table S1.

$I_{CaL}$ current and $J_{up}$ flux were reduced to better match the experimental data. Maintaining such levels of current augmentation in the ToR-ORd model was not possible since the model started developing a high number of repolarization abnormalities in basal populations (no β-ARS). This arrhythmogenic behaviour was caused by the different formulation of some of the currents in the ToR-ORd model, that present higher values than the ORd version. Some of these main currents (especially the $I_{CaL}$ current) can be observed in Supplemental Figure 3, where we compared our model results with the ones given by the model developed by Gong et al [6]. The reduction of the $I_{CaL}$ current was also motivated by the new experimental results (Figure 2 of the main manuscript), which presented similar maximum $I_{CaL}$ values, far from the 40% difference reported in previous works [4].

| Ionic Current | HCM-ORd | HCM-ToR-ORd | Ionic Current | HCM-ORd | HCM-ToR-ORd |
|---|---|---|---|---|---|
| $I_{Na}$ | - | - | $I_{to}$ | -70% | -70% |
| $I_{NaL}$ | +165% | +165% | $I_{K1}$ | -30% | -30% |
| $I_{Nab}$ | +165% | +165% | $I_{Kr}$ | -45% | -45% |
| $I_{CaL}$ | +40% | +10% | $I_{Ks}$ | -45% | -45% |
| $\tau$ $I_{CaL}$ *fast inactivation* | +35% | +35% | $I_{NaK}$ | -30% | -30% |
| $\tau$ $I_{CaL}$ *slow inactivation* | +20% | +20% | $J_{Up}$ | -25% | -35% |
| $I_{NCX}$ | +30% | +30% | $J_{Rel}$ | -20% | -20% |

**Table S1.** Comparison of the ionic electrophysiological remodelling in HCM-ORd vs HCM-ToR-ORd.

Most of the previous modifications were driven by intracellular calcium handling abnormalities characteristic of HCM patients[7][8]. As informed by the experimental data, the characteristic long action potential duration (APD) of HCM patients can also be an indicator of a less functional calcium dependant inactivation (CDI) of the $I_{CaL}$ channels. In healthy myocytes, an augmentation in the magnitude of $I_{CaL}$ currents is accompanied by an increase in the CDI and therefore a reduction of the APD. As stated in Table S1, in HCM cells $I_{CaL}$ currents are upregulated in comparison to healthy myocytes, therefore we modified the CDI in our HCM model to revert this behaviour and obtain longer APDs. In particular, we added a voltage shift in the $j_{ca}$ gate:

$$jca_{\infty} = \frac{1}{1 + \exp\left(\frac{(V + 18.08 + 12.0)}{2.7916}\right)}$$

Thanks to the previous modification, the HCM model is more resistant to developing repolarization abnormalities. Otherwise, the model was not able to reproduce long APDs, presenting EADs in almost all the cases where a certain APD (around 450ms) was surpassed, especially under ISO administration.

### Modifications to the β-ARS cascade (control population)

The different formulation of the ToR-ORd model also affected the implementation of the β-ARS phosphorylation cascade. We used the Heijman model[9], adapted to human (ORd model) in different works [6,10]. However, to replicate experimental results and obtain realistic electrophysiological outcomes, several changes needed to be implemented, especially in $I_{CaL}$ and $I_{Ks}$ channels:

$I_{CaL}$ values were fitted against human experimental data published by Magyar et al. [11] (used in the calibration of the ToR-ORd original model), Chen et al. [12] and canine $I_{CaL}$ data from Farkas et al. [13]. We maintained the original I-V, activation and inactivation curves of the base control model (no β-ARS) to preserve the behaviour of the original ToR-ORd model. These curves were calibrated using the data of Magyar et al. [11]. Unfortunately, there was no data regarding β-ARS in the same study, so we adapted the ISO response from other studies.

Following the same procedure in Tomek et al. [1] (Appendix 3), we obtained the $I_{CaL}$ activation by dividing the I-V curve reported in [13] by the Goldman-Hodgkin-Katz driving force, computed using Davies equation and fitted the activation curve under ISO application using a Gompertz function. Since Magyar's activation curve (Fig S1.B, black) is shifted with respect to the Farkas' activation curve (solid green line), we applied a voltage shift in the calculated curve to maintain the shift of the β-ARS curve with respect to control. The resulting activation curve, represented if Fig S1.A is given by the expression:

$$d_{\infty,PKA} = 1.0323 \cdot \exp\left(-1.0553 \cdot \exp\left(-0.0810 \cdot (V + 9.5)\right)\right)$$

Farkas' experimental procedure and activation curve in control were more similar to the ones in Magyar experiment and allowed a more straightforward adaptation of the activation curve under β-ARS. A comparison of the different experimental curves can be seen in Fig S1.B. The obtained activation curve under β-ARS presents a shift of -9.5 mV with respect to the activation curve in control, similar to the shift reported in both experimental works.

I-V curves were also computed and are represented in Fig. S1.C. The increase of the $I_{CaL}$ maximal current under PKA phosphorylation was slightly reduced to match the augmentation reported by human experimental data and to reproduce the observed APD shortening under β-ARS:

$$PCa_{PKA} = PCa \cdot 1.1$$

A comparison between the normalised obtained curves and normalised experimental data can be seen in Fig. S1.D. It can be observed that the gain that the I-V curve presents in the developed model under β-ARS is at the same level as the one presented in human data (Chen et al). The gain obtained in canine data (Farkas et al.) is considerably higher (close to 2,5 fold). Other studies with canine data have also reported similar I-V curve gain [14][15]. Finally, $I_{CaL}$ inactivation curve under β-ARS was also modified by applying a similar shift like the one observed in Farkas et al. (Fig S1.E).

Potassium currents are increased during β-ARS effect. However, due to the smaller basal $I_{Ks}$ values of the ToR-ORd model, the increase in $I_{Ks}$ could not reach the values required for obtaining a shortening of the APD. Therefore, following a similar approach reported in Gong et al., we increased the conductance gain of the channel to match experimental observation. This increase is also motivated by several electrophysiological studies that reported a huge increase in the $I_{Ks}$ under β-ARS [16][17].

$$GKs_{PKA} = GKs \cdot 50$$

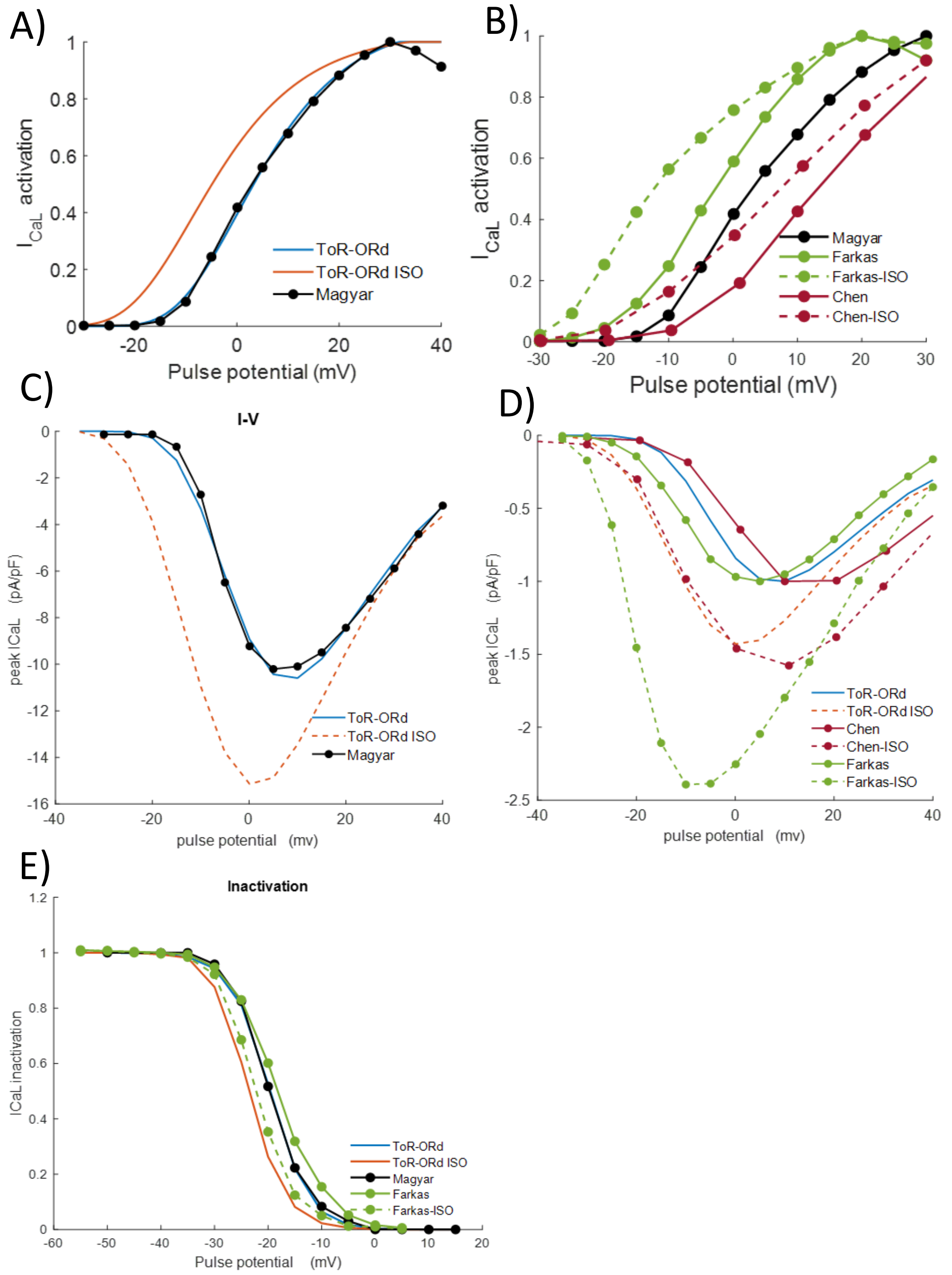


*Figure S1: A) $I_{CaL}$ activation curves of the model under β-ARS B) Comparison of the different experimental activation curves used in the calibration of the model C) Obtained I-V relationship of the model under β-ARS D) Comparison between the model I-V curves and different experimental I-V curves. E) Comparison between the model $I_{CaL}$ inactivation and experimental inactivation curves.*

The effects of the BARS in all the phosphorylated currents are listed below:

**ICaL**

$$PCa_{PKA} = PCa \cdot 1.1$$

$$d_{\infty,PKA} = 1.0323 \cdot \exp\left(-1.0553 \cdot \exp\left(-0.0810 \cdot (V + 9.5)\right)\right)$$

$$f_{\infty,PKA} = \frac{1}{1 + \exp\left(\frac{(V + 19.58 + 4.0)}{3.696}\right)}$$

**INa**

$$GNa_{PKA} = GNa \cdot 1.7$$

$$h_{\infty,PKA} = \frac{1}{\left(1 + \exp\left(\frac{(V + 71.55 + 5.0)}{7.43}\right)\right)^2}$$

$$h_{\infty,CaMK,PKA} = \frac{1}{\left(1 + \exp\left(\frac{(V + 71.55 + 6 + 5.0)}{7.43}\right)\right)^2}$$

**IKs**

$$GKs_{PKA} = GKs \cdot 50$$

$$x_{s1,\infty,PKA} = x_{s1,\infty}$$

$$x_{s2,\infty,PKA} = x_{s1,\infty,PKA}$$

$$\tau_{xs1,PKA} = -263{,}4 + 817.3 + 2.75 \cdot \cdot \frac{1}{2.326 \cdot 10^{-4} \cdot \exp\left(\frac{V + 48.28}{17.80}\right) + 1.292 \cdot 10^{-3} \cdot \exp\left(\frac{-(V + 210.0)}{230.0}\right)}$$

$$\tau_{xs2,PKA} = \tau_{xs1,PKA}$$

**INaK**

$$K_{Nai,PKA} = 0.7 \cdot K_{Nai}$$

**IKb**

$$G_{Kb,PKA} = 1.2 \cdot G_{Kb}$$

**Jrel**

$$\alpha_{rel,PKA} = 1.4 \cdot \alpha_{rel}$$

$$\tau_{rel,PKA} = 0.75 \cdot \tau_{rel}$$

$$\alpha_{rel,CaMK,PKA} = 1.4 \cdot \alpha_{rel,CaMK}$$

$$\tau_{rel,CaMK,PKA} = 0.75 \cdot \tau_{rel,CaMK}$$

**Jup**

$$J_{up,NP,PKA} = \frac{0.005425 \cdot [\mathrm{Ca}^{2+}]_i}{([\mathrm{Ca}^{2+}]_i + 0.00092 \cdot 0.7)}$$

$$J_{up,CaMK,PKA} = 2.75 \cdot \frac{0.005425 \cdot [\mathrm{Ca}^{2+}]_i}{[\mathrm{Ca}^{2+}]_i + (0.00092 - 0.00017) \cdot 0.7}$$

**TnI**

$$K_{m,TRPN,PKA} = 1.6 \cdot K_{m,TRPN}$$

### Modifications to the β-ARS cascade (HCM population)

Experimental observations in HCM cells under β-ARS showed that these cells have an altered β-ARS response. In particular, we observed a reduction in the gain of the potassium currents and a slower $I_{CaL}$ inactivation. We applied the following changes in the currents based on these data:

**ICaL**

$$\tau_{f,fast,HCM} = 2 \cdot \tau_{f,fast}$$

$$\tau_{f,slow,HCM} = 2 \cdot \tau_{f,slow}$$

$$\tau_{f,Ca,fast,HCM} = 2 \cdot \tau_{f,Ca,fast}$$

$$\tau_{f,Ca,slow,HCM} = 2 \cdot \tau_{f,Ca,slow}$$

**IKs**

$$GKs_{PKA} = GKs \cdot 5$$

**IKb**

$$G_{Kb,PKA} = 1.02 \cdot G_{Kb}$$

**EXTENDED RESULTS**

### Additional data on the electrophysiological response

Figure S2 shows with more detail the simulated representative calcium transient cases in control and HCM models. As reported experimentally [4], HCM calcium transient has a similar amplitude to the control population and a slower decay. Under β-ARS, the amplitude of CaT in control and HCM increases whereas both exhibit a faster decay.

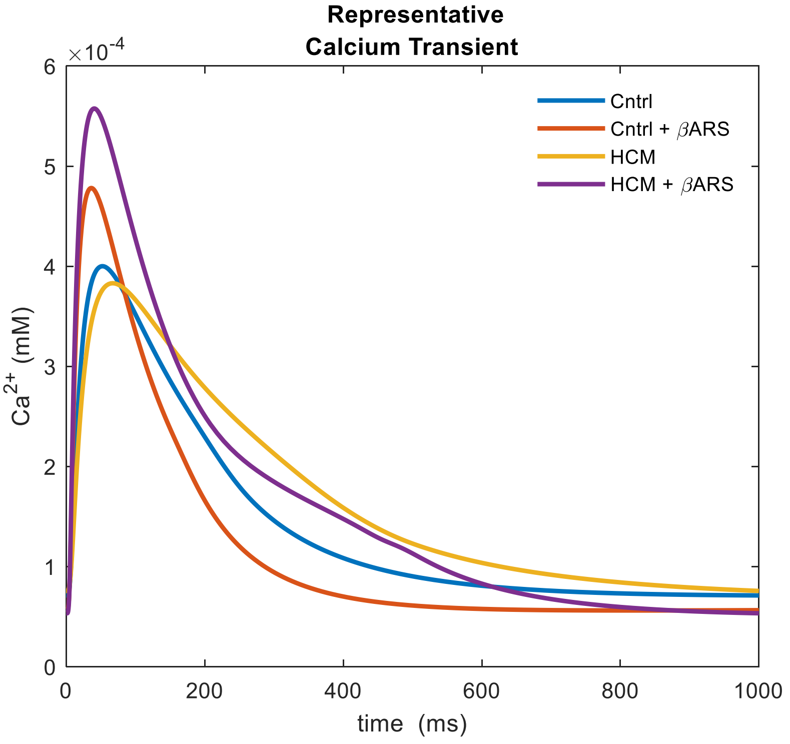


*Figure S2: Simulated calcium transients (CaT) in control and HCM baseline models at 1 Hz*

### Model comparison

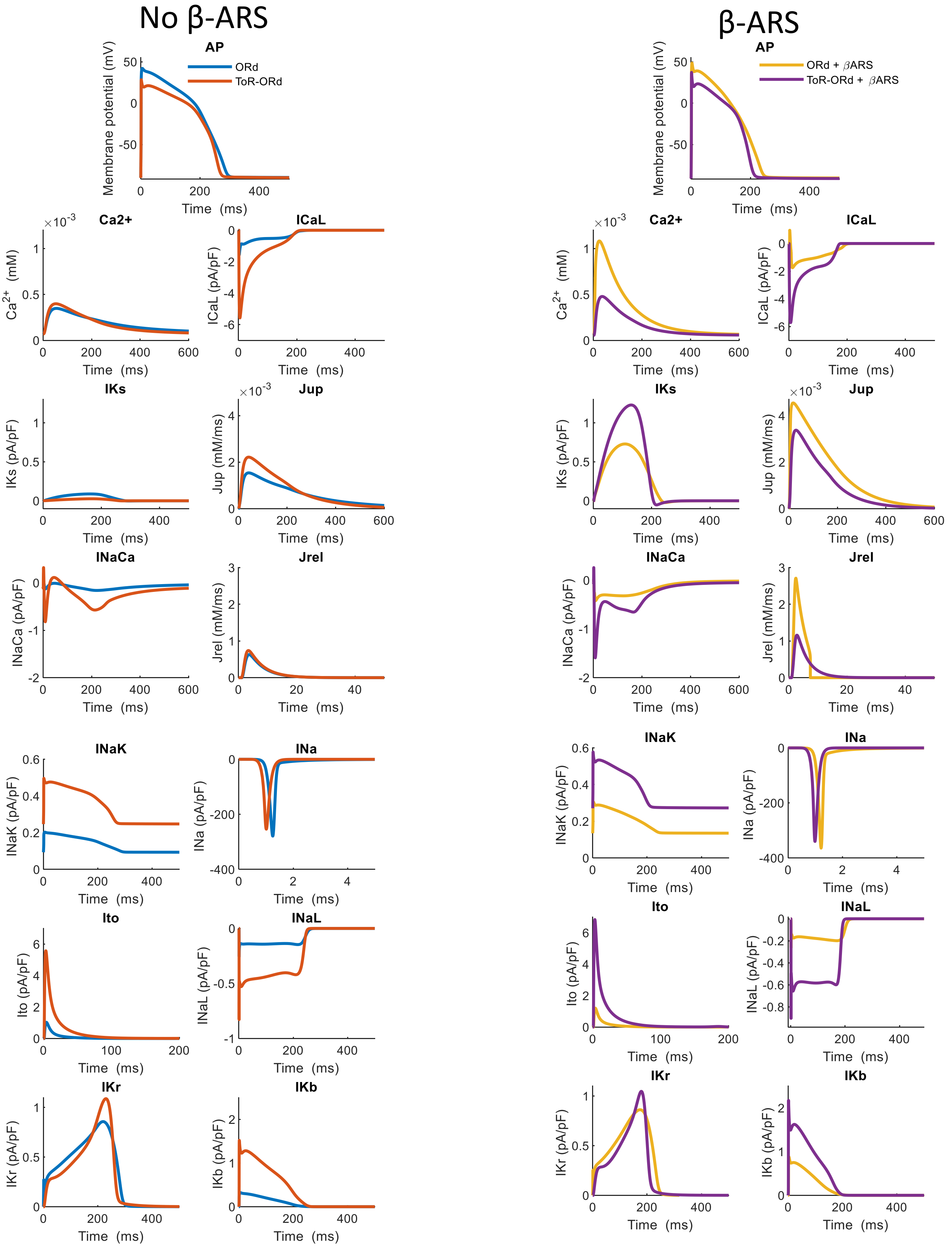


*Figure S3: Comparison between the different action potentials and ionic currents of the ORd model and ToR-ORd model, with and without the effect of β-ARS.*
